# Differential expression and function of SVIP in breast cancer cell lines and *in silico* analysis of its expression and prognostic potential in human breast cancer

**DOI:** 10.1101/2023.03.03.530947

**Authors:** Esra Atalay Şahar, Petek Ballar Kirmizibayrak

## Abstract

The heterogeneity of cancer strongly suggests the need to explore additional pathways to target. As cancer cells have increased proteotoxic stress, targeting proteotoxic stress-related pathways such as endoplasmic reticulum stress is attracting attention as new anticancer treatments. One of the downstream responses to endoplasmic reticulum stress is endoplasmic reticulum-associated degradation (ERAD), a major degradation pathway that facilitates proteasome-dependent degradation of unfolded or misfolded proteins. Recently, SVIP, an endogenous ERAD inhibitor, has been implicated in cancer progression, especially in glioma, prostate, and head and neck cancers. Here, the data of several RNA-seq and gene array studies were combined to evaluate the SVIP gene expression analysis on a variety of cancers, with a particular focus on breast cancer. SVIP was found to be overexpressed in primary breast tumors compared to normal tissues correlated well with its promoter methylation status and genetic alterations. Similarly, immunoblotting analysis showed that SVIP was expressed significantly higher in breast cancer cell lines compared to non-tumorigenic epithelial cell line. On the other hand, the expression of the key proteins of gp78-mediated ERAD did not exhibit such a pattern. Interestingly, silencing of SVIP enhanced the proliferation of p53 wt MCF7 cells but not p53 mutant T47D cells, however increased migration ability of both cell lines. Interestingly, SVIP expression is high in primary breast tumors but low in breast metastatic tumors. This correlates well with a lower probability of survival of breast cancer patients with lower SVIP expression compared to the patients with overexpressed SVIP. Overall, our data revealing the differential expression and function of SVIP on breast cancer cell lines together with *in silico* data analysis suggest that SVIP may have complex functions in breast cancer progression and has the potential to be a therapeutic target for breast cancer.

## 1. Introduction

Breast cancer is the most commonly diagnosed cancer in women. The incidence rate of breast cancer in women surpassed lung cancer, with 2.3 million new cases (11.7% of total cases) recorded in 2020, making it the most prevalent cancer type worldwide (Harbeck et al., 2019; Sung et al., 2020). Even though great efforts have been made to identify new therapeutic strategies or potential biomarkers by different approaches, breast cancer still remains the leading cause of cancer-related mortality in women. Given its heterogeneity with high rates of relapse and drug resistance, the identification of new prognostic biomarkers or therapeutic targets is still urgently needed in breast cancer research and treatment.

The endoplasmic reticulum is a central organelle that is involved in the maintenance of cellular homeostasis and the balance between health and disease. It regulates proper protein folding, modification and quality control (Avril et al., 2017; Lin et al., 2019). Multiple adverse external and intracellular factors, such as hypoxia, nutrient deprivation, cancer, obesity, neurodegeneration, oxidative stress, and viral infections, can disrupt the endoplasmic reticulum homeostasis. When endoplasmic reticulum homeostasis is disrupted and the protein folding capacity is exceeded, cells trigger a condition defined as “endoplasmic reticulum stress” (Jiang et al., 2020). The cellular response to endoplasmic reticulum stress is the unfolded protein response (UPR), a well-established signaling pathway. In order to reestablish normal functioning of endoplasmic reticulum and proteostasis, UPR activates several strategies in parallel and in series. These strategies include prevention of protein aggregation via enhancing endoplasmic reticulum chaperon expressions, attenuation of global protein translation, and promoting degradation of misfolded and unfolded proteins from the endoplasmic reticulum using the endoplasmic reticulum-associated degradation (ERAD) pathway (Sisinni et al., 2019; Wu et al., 2020; Jiang et al., 2020).

ERAD is a multi-component and complex system that involves the recognition of misfolded proteins by the appropriate luminal chaperones, retrotranslocation into the cytosol, polyubiquitination, and degradation by 26S proteasome (Tsai and Weissman, 2010; Segura-Cabrera et al., 2017). The AAA-ATPase Valosin-containing protein (VCP, also known as p97) is the key protein functioning in the retrotranslocation of ERAD substrates across the endoplasmic reticulum membrane into the cytosol. Besides ERAD, p97/VCP also participates in autophagy and mitophagy (Bento et al., 2018; Mengus et al., 2022). As it has also been revealed that p97/VCP is involved in cancer cell reprogramming (Sun and Qiu, 2020), the regulation of p97/VCP activity is crucial for cellular homeostasis. There are several identified functional partners of p97/VCP; three of them, namely E3 ubiquitin ligase gp78, small VCP/p97-interacting protein (SVIP), and VCP-interacting membrane protein (VIMP, also known as Selenoprotein S) possess VCP-interacting motif (VIM) in their structure. While gp78 and VIMP promote the ERAD process, SVIP acts as an endogenous ERAD inhibitor (Ballar et al., 2007; Ballar and Fang, 2008; Capelle et al., 2021).

Recent studies have suggested that SVIP may be involved in cancer progression. Firstly, SVIP levels were downregulated after androgen treatment in prostate cancer cells, while other components of ERAD machinery were upregulated (Erzurumlu and Ballar, 2017), which was found to be positively related to prostate tumorigenesis. Similarly, SVIP was reported to be downregulated by androgen in the glioma cells and suggested as new target for new for p53wt gliomas (Bao et al., 2017). In a study that aimed to identify putative genetic and epigenetic changes in the p97/VCP-mediated ERAD pathway in human tumors, SVIP promoter CpG island was found to be methylated in 50% (19 of 38) of head and neck cancer cell lines. Additionally, SVIP undergoes DNA hypermethylation also in esophageal (8 of 35, 23%) and cervical (3 of 14, 21%) cancers and hematological malignancies (22 of 154, 14%), particularly B cell lymphoma (15 of 45, 33%). Apart from these cancer types, the SVIP promoter CpG island was most often found to be unmethylated in the other cancer types (Llinàs-Arias et al., 2019). Moreover, comparison of SVIP expression in different prostate cell lines, revealed that SVIP is highly expressed in androgen-dependent prostate cancer cells (LNCAP, 22RV1) but not in androgen-independent cell lines (PC3, DU145) or non-tumorigenic prostate cell lines including normal prostate epithelial cell line (RWPE1) and benign prostatic hyperplasia epithelial cell line (BPH1) (Erzurumlu and Ballar, 2017).

Considering all this data suggesting that SVIP may play a role in tumor progression,, we have first performed *in silico* analysis of SVIP expression on various cancers, with a particular focus on breast cancer. We examined the association between SVIP and breast cancer by performing comprehensive bioinformatics analysis on several large online databases and evaluated the expression profile in variety of breast cancel cell line. While SVIP expression was much higher in breast cancer cell lines compared to nontumoral MCF10A cells, SVIP silencing augmented the proliferation of p53 wt MCF7 cells, but not p53 mutant T47D cells. Our data revealing the differential expression and function of SVIP on breast cancer cell lines together with *in silico* data suggest that SVIP may have complex functions in breast cancer progression and has the potential to be an important biomarker for clinical prediction of breast cancer.

## 2. Methods

### 2.1. Gene expression analysis

SVIP gene expression levels were analyzed through the TNMplot database (http://www.tnmplot.com) (Bartha and Győrffy, 2021). The Tumor Immune Estimation Resource (TIMER) database (https://timer.comp-genomics) (Li et al., 2020) was used to verify the expression of SVIP in various cancers. TNMplot database (http://www.tnmplot.com) with RNA-sequence data was used to explore the SVIP gene expression in breast cancer tumor, normal, and metastatic tissues. TNMplot database provides gene array data from the Gene Expression Omnibus of the National Center for Biotechnology Information (NCBI-GEO), RNA-seq datasets from The Cancer Genome Atlas (TCGA) and also Therapeutically Applicable Research to Generate Effective Treatments (TARGET) and The Genotype-Tissue Expression (GTEx).

### 2.2. Promoter methylation analysis of SVIP

With data obtained from TCGA project, UALCAN database is an interactive web portal which enables gene expression analysis on about 20.500 protein-coding genes in 33 different tumor types (Chandrashekar et al., 2017). In addition, this tool was also used to find the promoter methylation of SVIP in breast cancer. Furthermore, the correlation between methylation and expression was analyzed using cBioPortal. cBioPortal for Cancer Genomics (v3.7.27 http://www.cbioportal.org/) database. The portal contains data sets from published cancer studies, including the Cancer Cell Line Encyclopedia (CCLE) and the TCGA pipeline (Cerami et al. 2012; Gao et al. 2013).

### 2.3. Genomic analysis

cBioPortal for Cancer Genomics (v3.7.27 http://www.cbioportal.org/) is a powerful platform that is used for exploring, visualizing, and analyzing multidimensional cancer genomics data (Cerami et al. 2012; Gao et al. 2013). The genomics features of SVIP on breast cancer were analyzed via the cBioPortal database. The gene alteration frequency of SVIP using TCGA, PanCancer Atlas from 1084 breast cancer patients was analyzed. The alterations included amplification, deep deletion, mRNA expression, and truncating mutation. Mutation details, copy number alterations (CNA), and genomic alteration of SVIP were also taken into consideration in different breast cancer types. CNA, mutation details, and mRNA expression were generated from RNA-seq (log2) data with default settings and plotted with mRNA expression data using the cBioPortal database. The mRNA expression z-score threshold was ± 2 between the unaltered and altered patients.

### 2.4. SVIP gene expression based on intrinsic molecular subtype and clinicopathological dataanalysis

The Breast Cancer Gene-Expression Miner v4.8 (bc-GenExMiner v4.8), a mining tool, UALCAN, and the Gene Expression Profiling Interactive Analysis (GEPIA) (http://gepia.cancer-pku.cn/) databases were used evaluation of SVIP expression based on clinicopathological characteristics and intrinsic molecular subtype (PAM50 cancer subtypes) (Chandrashekar et al., 2017; Tang et., 2019; Jézéquel et al., 2021). The clinicopathological parameters were as follows: estrogen receptor status (ER), progesterone receptor status (PR), HER2 receptor status, p53 status, pathological tumor stage, and nature of the tissue (healthy, tumor, and tumor-adjacent). All data were from TCGA and GTEx datasets.

### 2.5. Survival analysis

The effects of SVIP expression on the survival of patients with breast cancer were performed using the Kaplan Meier Plotter online survival analysis tool (https://kmplot.com/analysis/) (Lanczky and Gyorffy, 2021). This tool is used in a meta-analysis-based discovery and validation of survival biomarkers. The prognostic indicators evaluated include overall survival (OS) and distant-metastasis-free survival (DMFS).

### 2.6. Cell culture and Treatments

All cell lines were grown as exponentially growing monolayers by culturing according to the American Type Culture Collection (ATCC, USA) instructions. MCF10A cells were grown in DMEM/F12 medium with 5% horse serum, 100 μg/ml EGF, 1 mg/ml hydrocortisone, 1 mg/ml cholera toxin, and 10 mg/ml insülin. MCF-7, SK-BR-3, MDA-MB-231 cells were cultured in Dulbecco’s modified Eagle’s High Glucose medium (DMEM) supplemented with 10% fetal bovine serum (FBS). BT474 cells were grown in DMEM High Glucose supplemented with 10% FBS, 10 μg/mL insülin, 1% MEM non-essential aminoacid. T47D cell line was grown in DMEM High Glucose supplemented with 10% FBS and 0.1% MEM non-essential aminoacid. ZR-75-1 cell line was grown in Roswell Park Memorial Institute medium (RPMI) 1640 containing 10% FBS. All cell lines were grown as monolayers and were incubated at 37 °C with 95% humidified air and 5% CO_2_. Cells were checked regularly for mycoplasma contamination by PCR (MycoAlert™ Mycoplasma Detection Kit, Lonza).

For experiments using 17β-estradiol (E2) (Cayman Chemical, USA), phenol red-free DMEM High glucose and Dextran-coated charcoal treated FBS (DCT-FBS) (Biological Industries, USA) were used. MCF7, T47D, ZR-75-1 (3×10^5^ cells/well) were cultured in six-well plates at a confluency of 40%; after 1 day, the normal medium was replaced by phenol red-free DMEM High glucose supplemented with 5% DCT -FBS. Before E2 treatments, cells were cultured for 72 h in phenol red-free medium supplemented with 5 % DCC-FBS. The medium was changed to 0.5% DCC-FBS at time 0 h., and E2 was added to a final concentration of 10 nM. E2 was then incubated for 30 m, 1 h, 2 h, 3h, and 4 h.

Transfections were performed with Lipofectamin-2000 (Invitrogen) following the manufacturer’s instructions, in order to manipulate protein expression levels either by silencing. Silencer® Negative Control siRNA (Ambion, 4611) and SVIP siRNA (AM16104, sense sequence: GACAAAAAGAGGCUGCAUC) were ordered from Ambion.

### 2.7. Immunoblotting

Cells were harvested and lysed using RIPA buffer (1X PBS, 1% nonidet P-40, 0.5% sodium deoxycholate, and 0.1% SDS, pH 8.0). The total protein concentrations were determined using the bicinchoninic acid (BCA) protein assay (Thermo Fisher Scientific, USA). 40 μg of total cellular proteins were loaded to the gels after denatured in 4× Laemmli buffer (Bio-Rad) at 37 °C for 1 hour. Proteins were separated by sodium dodecyl sulfate–polyacrylamide gel electrophoresis (SDS-PAGE) and transferred to polyvinylidene fluoride (PVDF) membranes (EMD Millipore; Thermo Fisher Scientific). PVDF membrane was treated with primary and secondary antibodies, and then, proteins were visualized using Clarity ECL substrate solution (Bio-Rad) by Fusion-FX7 (Vilber Lourmat; Thermo Fisher Scientific, USA). β-Actin (Sigma-Aldrich, A5316) and GAPDH (CST-5174) were used as loading controls. The primary antibody for SVIP (HPA039807) was purchased from Sigma-Aldrich. The antibodies against gp78 (9590), Hrd1 (147773), Derlin1 (8897), ERα (8644) were purchased from Cell Signaling Technology and p97/VCP (612182) was obtained from BD Transduction Laboratories. Secondary antibodies (Goat anti-rabbit-31460 and Goat anti-mouse-31430) were purchased from Thermo Fisher Scientific. All western blot experiments were performed at least in three independent replicates.

### 2.8. Cell proliferation and migration

Proliferation rate of SVIP siRNA-transfected and negative siRNA transfected MCF7 and T47D cells were monitored using the xCELLigence impedance-based real-time cell analysis system (ACEA Biosciences, USA). MCF7 and T47D cells were seeded (8000 cells/well) on an E-plate-16 at the optimal cell density and cell proliferation was monitored every 30 min. The electrical impedance measured by the RTCA software and data were represented as cell index. Impedance is correlated with an increase in the number of cells on well by measuring cell index.

The wound healing assay SVIP siRNA-transfected and negative siRNA transfected MCF7 and T47D cells were plated at 2 ×10^5^ cells per well into Cytoselect wound-healing assay 24 well plate (Cell Biolab, Inc). Each well contained an insert that created a wound field. After 24 h, the insert was gently removed creating a gap of 0,9 mm. The migration of the cells into the wound field was monitored for 72h and images were taken with microscope (Olympus CKX41). The analysis of wound closure % was determined by using the ImageJ software (http://imagej.nih.gov/ij/).

The migratory ability of MCF7 and T47D cells transfected with SVIP siRNA was assessed using 24-well Transwell inserts (8-μm pore size; Greiner Bio-one, Austria). A total of 1×10^4^ cells were plated in the upper chambers of Transwell filters in 100 μl phenol red free DMEM with 5% DCT-FBS. The cell migration was stimulated through the membranes by DMEM (650 μl) containing 20% FBS in the lower chambers as a chemoattractant. Subsequent to 48 h of incubation at 37°C, the migratory cells on the lower membrane surface of the insert were fixed with methanol at room temperature and stained with 0.05% crystal violet solution (Serva). Migration was quantified by counting stained cells in five randomly selected fields using microscope and the data were expressed as the mean percentage of migrated cells compared with the vehicle control groups.

### 2.9. Statistical analysis

The significant differences of SVIP expression level in the different cancer tissues were analyzed by the Mann-Whitney U test in the TNM plot database (* *p < 0*.*01*). Then, distributions of SVIP gene expression levels in the different cancer type was computed by the Wilcoxon test in the TIMER database (*: *p < 0*.*05*; **: *p <0*.*01*; ***: *p <0*.*001*). The tumor, normal and metastatic breast cancer data were compared using Kruskal–Wallis test in TNM plot database. The statistical significance cutoff was set at *p < 0*.*01*. In Kaplan–Meier Plotter database the survival rate with *p* values was analyzed by Log-Rank test. *p < 0*.*05* was considered statistically significant. The cutoff value of SVIP gene expression was chosen as median. Dataset was split into two groups of patients and plots generated accordingly. Welch’s T-test and Dunnett–Tukey–Kramer’s tests estimated the significance of SVIP expression levels based on clinicopathological features in the bc-GenExMiner v4.8 (*p <0*.*001*). The differences in SVIP promoter methylation levels between breast cancer and normal tissue were analyzed by Welch’s T-test in the UALCAN database. *p < 0*.*05* was considered significant. Data are presented as means ± standard deviation (SD). For experiments comparing differences between groups were performed by Student’s t-test using GraphPad Prism software. The significance threshold was accepted as *p < 0*.*05*. The correlation between SVIP expression and methylation was evaluated using Spearman’s and Pearson correlation analysis and statistical significance in cBioPortal. The correlation of gene expression was evaluated using Spearman’s correlation and the p-value in TIMER. *p < 0*.*05* was considered statistically significant if not especially noted.

## 3. Results

### 3.1. The expression of SVIP in Different Cancer Types

To investigate possible differential expression of SVIP in tumor and normal tissues, the gene expression of SVIP was compared in normal and tumor tissues using the TNMplot database. The expression of SVIP in tumor tissue was significantly higher than in normal tissue in most cancer types, including acute myeloid leukemia, breast, colon, liver, pancreas, thyroid, and prostate cancers (**Fig. 1A**). Interestingly, SVIP expression was higher in renal chromophobe cell carcinoma (Renal CH) and lung adenocarciona (Lung AC), while lower in renal clear cell carcinoma (Renal CC), renal papillary cell carcinoma (Renal PA), and lung squamous cell carcinoma (Lung SC) than in normal tissue (Fig. 1A). When TIMER database with RNA-seq data of multiple malignancies in TCGA was used to verify the expression of SVIP in pan-cancer, the analysis revealed that breast invasive carcinoma (BRCA), cholangiocarcinoma (CHOL), colon adenocarcinoma(COAD), liver hepatocellular carcinoma (LIHC) and lung adenocarcinoma (LUAD) had higher SVIP expression, while glioblastoma multiforme (GBM), head and neck squamous cell carcinoma (HNSC), kidney renal clear cell carcinoma (KIRC), kidney renal papillary cell carcinoma (KIRP), and lung squamous cell carcinoma (LUSC) had lower expression of SVIP than in normal tissue. Importantly, the expression of SVIP was significantly upregulated in all the subtypes of breast cancer, namely luminal A, luminal B, HER-2 positive, and triple-negative (basal-like) types (**Fig. 1B**). Taken together, these results demonstrated that the SVIP gene expression was differentially regulated in most cancer types compared with the normal tissues, particularly in breast cancer (**Fig. 1A, 1B, S1**).

**Figure 1.**
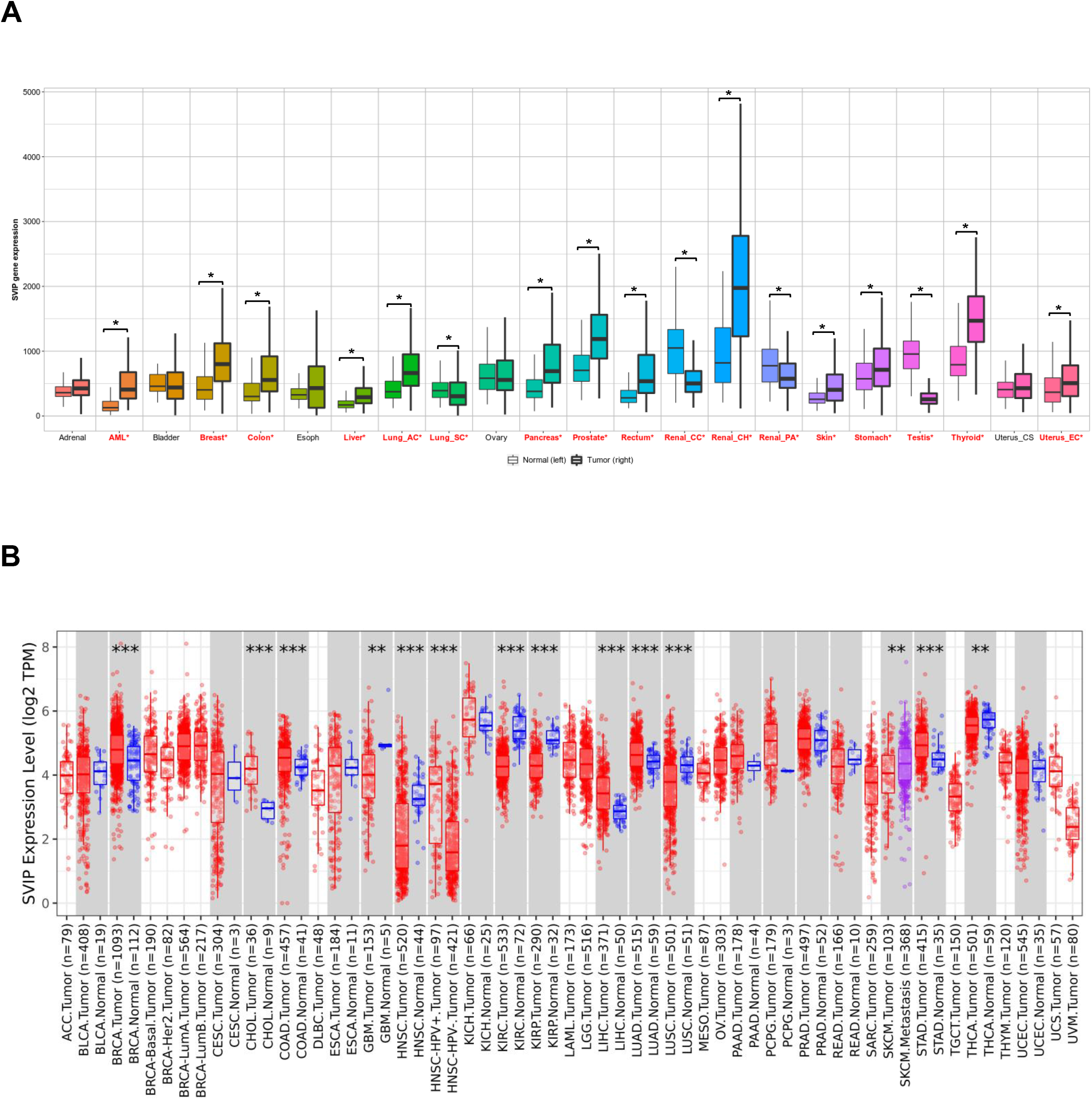
SVIP expression levels in different human tumors. **A)** Increased or decreased expression of SVIP in different tumors compared to normal tissues via the TNMplot database. Significant differences by Mann-Whitney U test are marked with red color (* *p < 0*.*01*). **B)** Human SVIP expression levels of different tumor types from the TCGA database was investigated by TIMER (* *p < 0*.*05*, ** *p < 0*.*01*, *** *p < 0*.*001*). Abbreviations: ACC: adrenocortical carcinoma; BLCA: bladder urothelial carcinoma; BRCA: breast invasive carcinoma; CESC: cervical squamous cell carcinoma; CHOL: cholangiocarcinoma; COAD: colon adenocarcinoma; DLBC: lymphoid neoplasm diffuse large B cell lymphoma; ESCA: esophageal carcinoma; GBM: glioblastoma multiforme; LGG: brain lower grade glioma; HNSC: head and neck squamous cell carcinoma; KICH: kidney chromophobe; KIRC: kidney renal clear cell carcinoma; KIRP: kidney renal papillary cell carcinoma; LAML: acute myeloid leukemia; LIHC: liver hepatocellular carcinoma; LUAD: lung adenocarcinoma; LUSC: lung squamous cell carcinoma; MESO: mesothelioma; OV: ovarian serous cystadenocarcinoma; PAAD: pancreatic adenocarcinoma; PCPG: pheochromocytoma and paraganglioma; PRAD: prostate adenocarcinoma; READ: rectum adenocarcinoma; SARC: sarcoma; SKCM: skin cutaneous melanoma; STAD: stomach adenocarcinoma; TGCT: testicular germ cell tumors; THCA: thyroid carcinoma; THYM: thymoma; UCEC: uterine corpus endometrial carcinoma; UCS: uterine carcinosarcoma; and UVM: uveal melanoma; AML: Acute myeloid leukemia; Lung_AC: lung adenocarcinoma; Lung_SC: lung squamous cell carcinoma; Renal_CC: renal clear cell carcinoma; Renal_CH: renal chromophobe cell carcinoma; Renal_PA: renal papillary cell carcinoma; Uterus_CS: uterine carcinosarcoma; Uterus_EC: uterine corpus endometrial carcinoma.

### 3.2. Promoter methylation level and genomic alteration analysis of SVIP in breast cancer

To investigate the possible relationship between SVIP expression and DNA methylation, the promoter methylation level of the SVIP in breast cancer tissues was analyzed using UALCAN tool from TCGA dataset. The level of methylation on the SVIP promoter was significantly lower in breast cancer compared to the normal tissues, as shown in **Fig. 2A** (*p*= 3.02E-04). Similar data was observed using cBioPortal database, where SVIP mRNA expression was negatively correlated with SVIP methylation (Spearman=-0.19, *p*=8.81e-7; Pearson=-0.10, *p*=0.0128) (**Fig. S2**).

**Figure 2.**
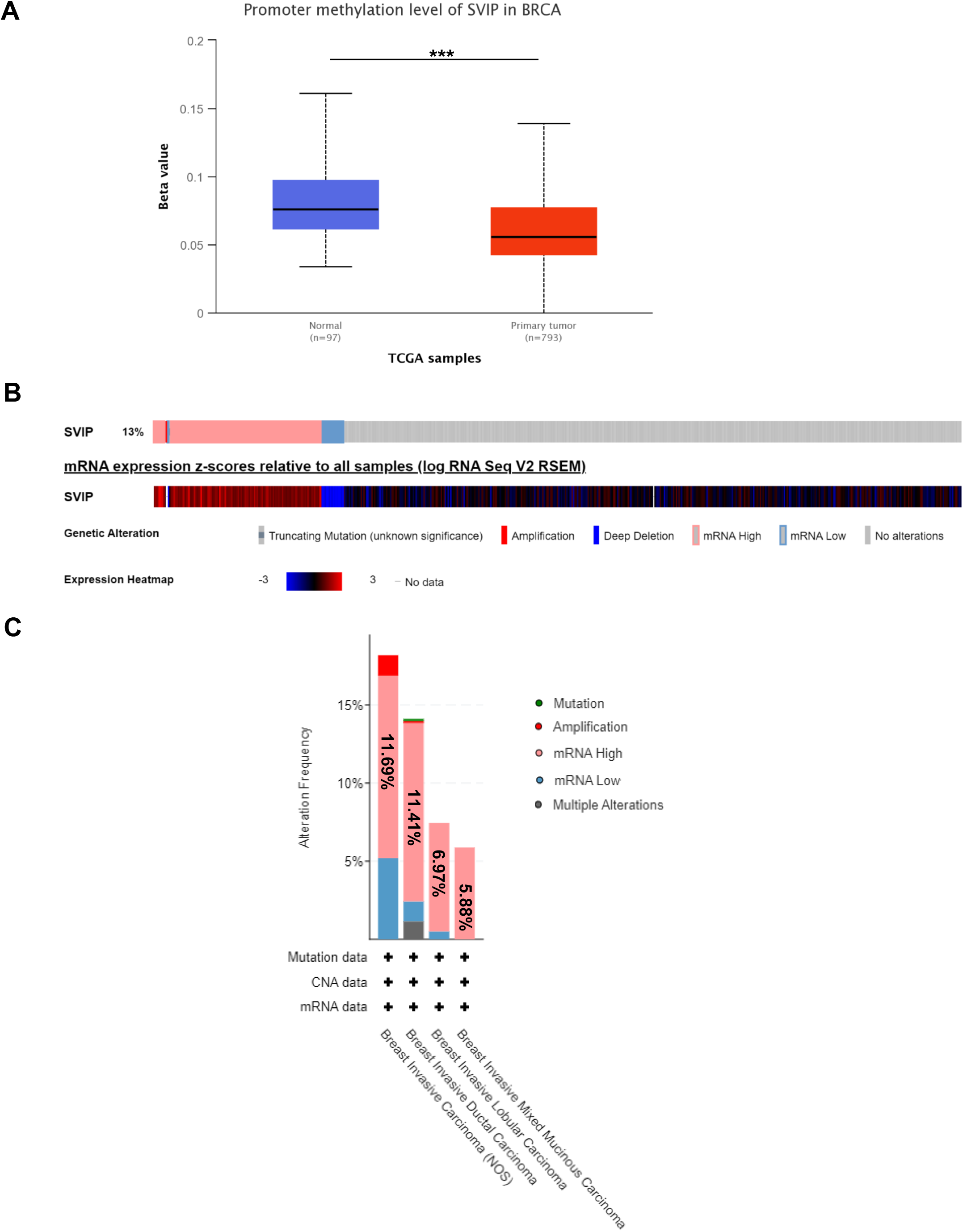
Promoter methylation level and genomic alterations of SVIP **A)** Promoter methylation level of SVIP in normal and breast cancer tissues (UALCAN) (p= 3.02E-04) ***, p < 0.001. **B)** Oncoprint of SVIP alteration in breast cancer (c-BioPortal). **C)** SVIP gene expression and mutation analysis in different breast cancer types (c-BioPortal).

We then explored the potential genetic alterations of SVIP in the context of breast cancer using the c-BioPortal online tool. The OncoPrint results showed in 142 of 1084 breast cancer patients (13%), the SVIP gene was altered either amplification, deep deletion, high mRNA expression or low mRNA expression. Among these alterations, the most common type was found to be high mRNA expression (**Fig. 2B**). When alteration frequency was assessed for different breast carcinoma samples, namely breast invasive carcinoma, breast invasive ductal carcinoma, breast invasive lobular carcinoma, and breast invasive mixed mucinous carcinoma, the frequency of SVIP gene alteration rates were varied from 5.88% to 11.69%, where mRNA high alteration was the most common one with the ratios varied from 5.88% to 18.18% (**Fig. 2C**).

### 3.3. Associations between SVIP expression levels and the clinicopathological parameters and its prognostic value in breast cancer patients

As SVIP is overexpressed in primary breast tumors compared to normal tissues, we next analyzed the expression of SVIP in distinct clinicopathological stages, parameters, and intrinsic subtypes of breast cancer. Firstly, SVIP was found to be highly expressed in every clinicopathological stages of breast cancer compared with normal tissues using the UALCAN portal (**Fig. 3A**). To further evaluate the relationship between SVIP expression levels and various clinicopathological parameters, the bc-GenExMiner (v4.8) online tool was utilized.

**Figure 3.**
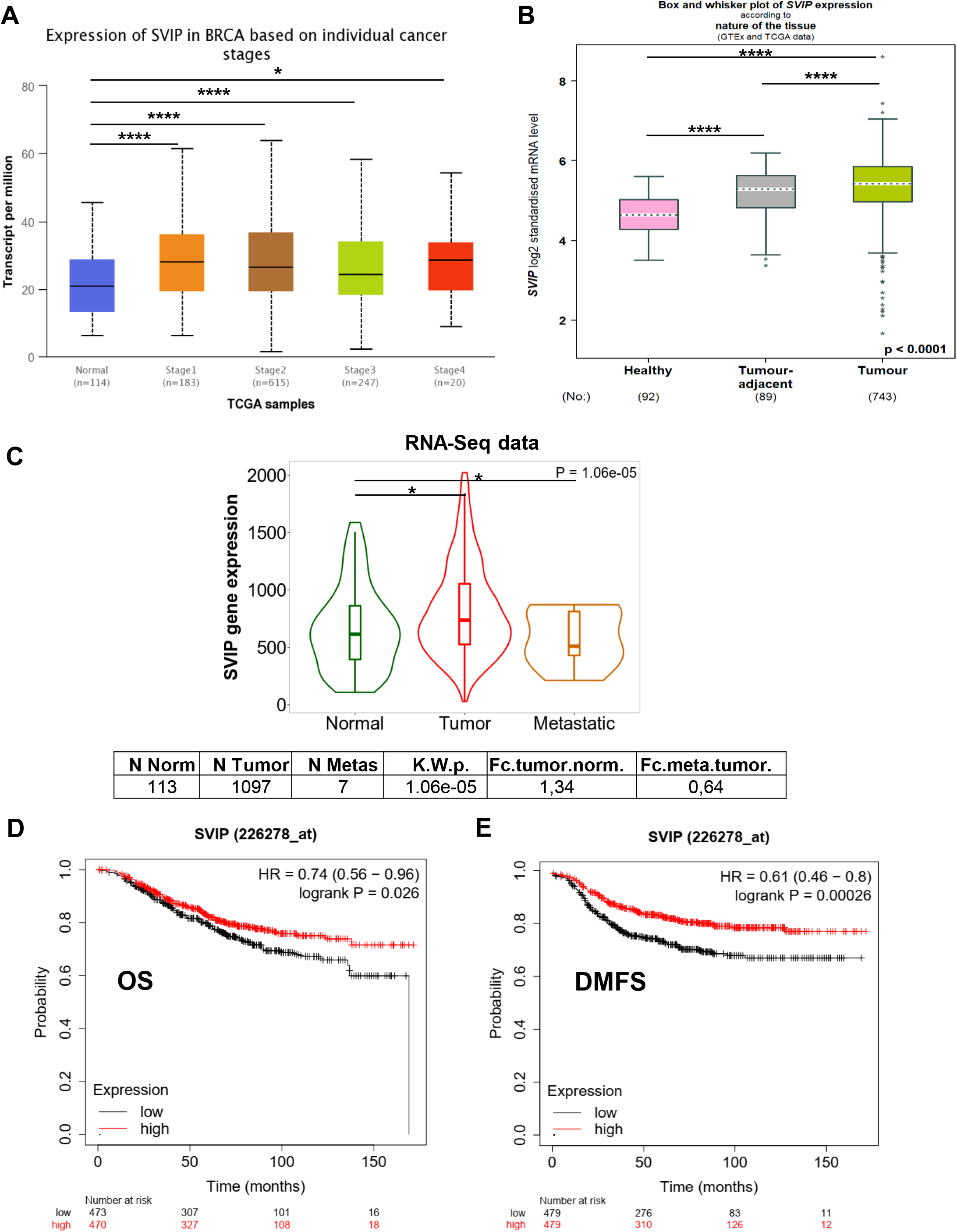
UALCAN, TNMplot, and bc-GenExMiner v4.8 portals analysis of breast cancer samples from the TCGA and GTEx datasets **A)** Expression of SVIP in different stages of breast cancer (UALCAN) (*, *p* < 0.05; **, *p* <0.01; ***, *p* < 0.001; ****, *p* < 0.0001). **B)** SVIP expression in breast cancer, tumor-adjacent normal tissue and healthy tissues (****, *p* < 0.0001) (bc-GenExMiner software). **C)** RNA-seq data of SVIP expression in normal, tumor, metastatic tissues by TNMplot database. **D-E)** Overall survival (OS) and Distant metastasis-free survival (DMFS) curves of BRCA by different expression levels of SVIP in the Kaplan-Meier Plotter database (*p* < 0.05).

SVIP was found to be highly expressed not only in the breast cancer tissues but also in the tumor-adjacent tissue compared to normal tissues (**Fig. 3B**). Intriguingly, higher SVIP mRNA levels were found in breast cancer patients with ER (+) than ER (−) and patients with PR (+) than PR (−) (**Fig. S3A, B**). On the other hand, SVIP mRNA levels were significantly lower in the HER2-positive group than in the HER2-negative group (**Fig. S3C**). The p53 wild-type status (sequence-based) was positively correlated with SVIP mRNA expression level (**Fig. S 3D**).

When the RNA-seq data of SVIP expression in normal, tumor, and metastatic breast tissues was evaluated via the TNMplot database, the analysis revealed a higher expression level of SVIP in tumor samples not only than in normal samples, but also in metastic samples (**Fig. 3C**). Interestingly, metastatic samples has lower SVIP expression compared to normal samples (*p=*1.06E-05).

Kaplan–Meier Plotter database was used to investigate the prognostic potential of SVIP in breast cancer patients. The RNA-seq data in Kaplan–Meier Plotter database, mainly extracted from Gene Expression Omnibus (GEO), European Genome-phenome Atlas (EGA), and TCGA. All results are displayed with *p-values* from a long-rank test. Results revealed that lower SVIP expression levels were associated with worse prognosis of overall survival (OS) (HR=0.74, *p*= 0.026) (**Fig. 3D**) and distant metastasis-free survival (DMFS) (HR=0.61, *p*= 0.00026) (**Fig. 3E**) in breast cancer. This result may be a reflection of decreased SVIP expression in metastatic breast tissues.

### 3.4. Expression of ERAD proteins in breast cancer cell lines

All these *in silico* observations prompted us to evaluate the SVIP protein expression levels by immunoblotting in model cell lines related to these breast cancer subtypes (**Fig. 4A**). For this aim, SVIP protein expression levels were analyzed in the cell lines that represent models for Luminal A [MCF7 (ER+, PR+, HER2-), T47D (ER+, PR+, HER2-) and ZR75-1(ER+, PR+/-, HER2-)], Luminal B [BT-474 (ER+, PR+, HER2+)], HER2+ [SKBR3 (ER-, PR-, HER2+)],

**Figure 4.**
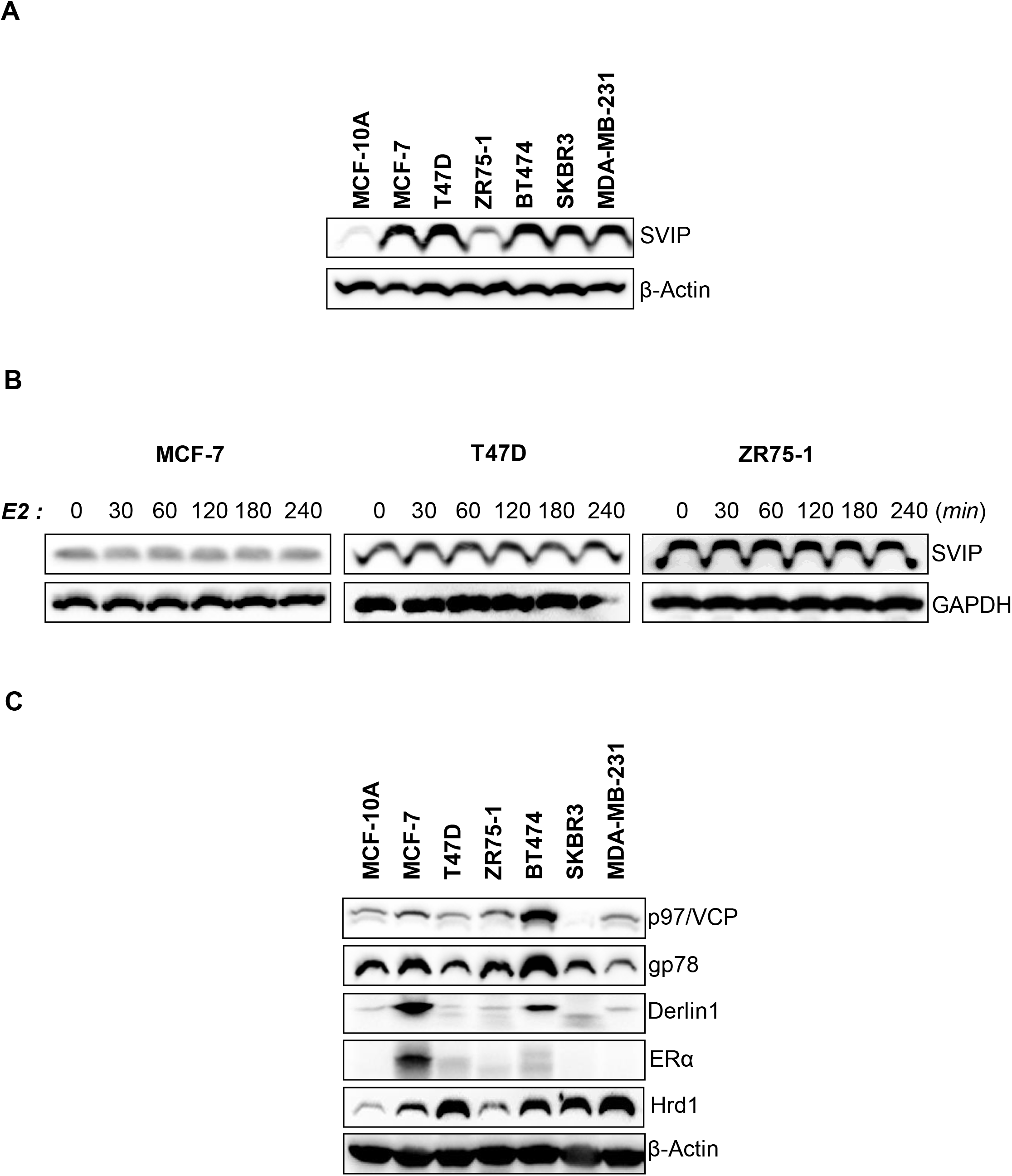
Expression of SVIP level in different breast cancer cell lines. **A)** SVIP expression in breast cancer cell lines and non-tumorigenic breast epithelial cell line by western blot analysis. Western blot is representative of three replicates with similar results. **B)** SVIP protein levels in breast cancer cell lines (MCF7, T47D, ZR-75-1) were determined by western blot analysis. β-actin or GAPDH was used as a loading control (n=3). **C)** Expression of ERAD components in different breast cancer cell lines.

Triple Negative Breast Cancer [MDA-MB-231 (ER-, PR-, HER2-)] breast cancer subtypes and non-tumorigenic breast epithelial cell line [MCF-10A ER-, PR-, HER2-] (Dai et al., 2017; Chen et al., 2020). Consisting with *in silico* data, SVIP expression was significantly higher in breast cancer cell lines compared to non-tumoral breast epithelial cell line (**Fig. 4A**). We have previously reported androgen dependent regulation of SVIP in prostate cancer (Erzurumlu and Ballar, 2017). Therefore, next, the estrogen dependency of SVIP expression in ER+ breast cancer cells was evaluated using 17β-estradiol (E2), and our results revealed that E2 did not cause any significant alterations in SVIP expression levels in MCF7, T47D, and ZR75-1 cells (**Fig. 4B**).

SVIP, the first identified endogenous ERAD inhibitor, inhibits gp78-mediated ERAD by competing with p97/VCP and Derlin1 (Ballar et al., 2007). Given that SVIP expression is high in breast cancer cells, we evaluated the other functional key players of gp78-mediated ERAD. The expression of p97/VCP, the key protein functioning on the retrotranslocation step of ERAD, is significantly higher in Luminal B subtype BT-474 cells but low in HER2+ subtype SKBR3 cells. gp78 and Derlin1 expression was higher in MCF7 and BT474 cells (**Fig. 4C**). Our results clearly indicated that breast cancer cell lines present significantly higher SVIP expression compared to non-tumorigenic epithelial MCF-10A cell line. On the other hand, the expression of other major proteins functioning in gp78-mediated ERAD, was high only in MCF7 and BT474 cells. Interestingly, the Hrd1 level, the other major ER resident ubiquitin ligase other than gp78, is also displayed expression pattern similar to SVIP expression.

### 3.5. The effect of SVIP in the proliferation and migration of breast cancer cell lines

Given the potential for the SVIP gene to be effective in breast cancer progression, functional analysis was performed in the next step. Firstly, the effect of SVIP on the proliferation of MCF7 and T47D cells was determined with the xCELLigence Real Time Cell Analyzer. Silencing of SVIP expression using siRNA oligonucleotides against SVIP mRNA enhanced the proliferation of MCF7, but not T47D cells (**Fig. 5A, B**).

**Figure 5.**
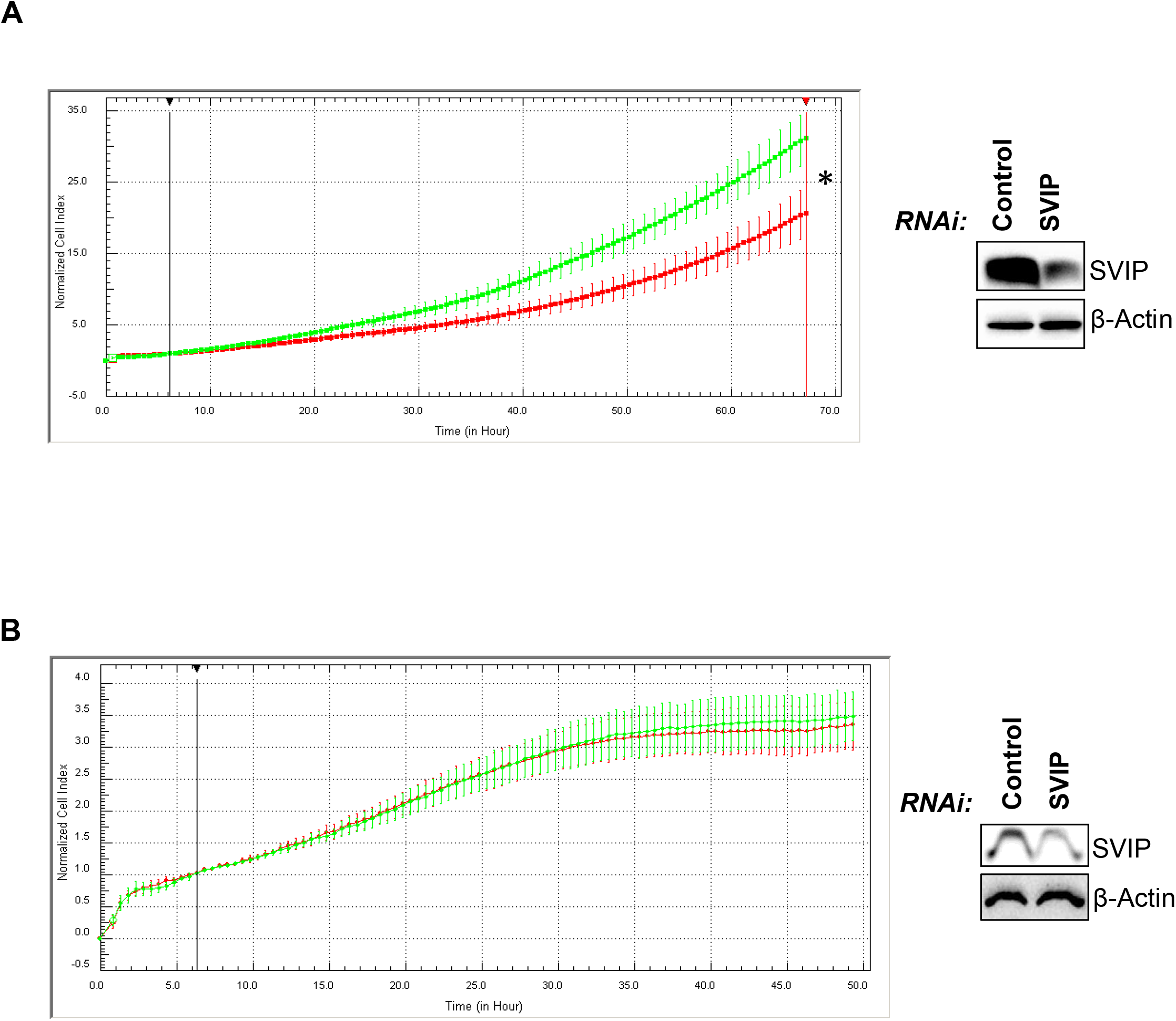
Effect of SVIP silencing on breast cancer tumorigenesis.**A)** Real-time proliferation analysis of MCF7 and **B)** T47D cells with xCELLigence RTCA. Cells were monitored for 48 h with three technical replicates (* p = 0.0003). MCF7 and T47D cells were transfected with siRNA oligonücleotides (SVIP and control siRNA). Immuno-blotting shows silencing of SVIP compared with MCF7 control cells (control siRNA). Error bars are presented as standard deviations.

Next, the role of SVIP silencing on the motility of MCF7 and T47D was tested by an *in vitro* wound healing model using Cytoselect 24 well plate which provides a standardized wound area via proprietary treated inserts. Our data indicated that silencing of SVIP increased the rate of wound closure of both MCF7 and T47D compared to that observed in negative siRNA-transfected control cells (*p < 0*.*05* for siSVIP MCF7, p < 0.05 for siSVIP T47D) (**Fig. 6A, 6B**). Consistingly, using Transwell Boyden chamber to analyse the migration ability revelead that silencing of SVIP increased the migration of both MCF7 and T47D cell lines (*p < 0*.*05* for siSVIP MCF7, *p < 0*.*05* for siSVIP T47D) (**Fig. 6C, 6D**). To conclude, our data suggest that SVIP has function on the regulation of the cell proliferation and migration of breast cancer cells.

**Figure 6.**
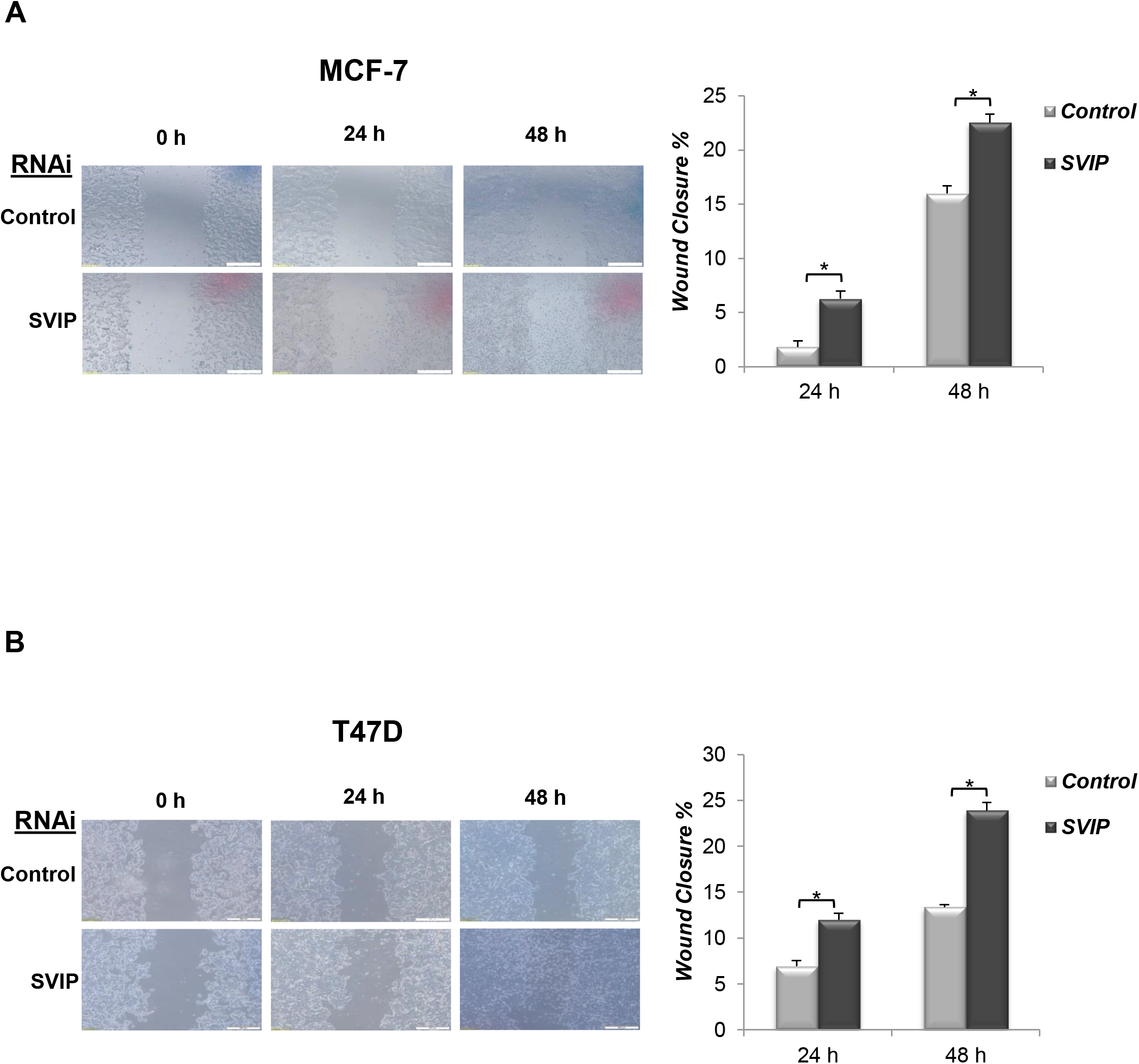
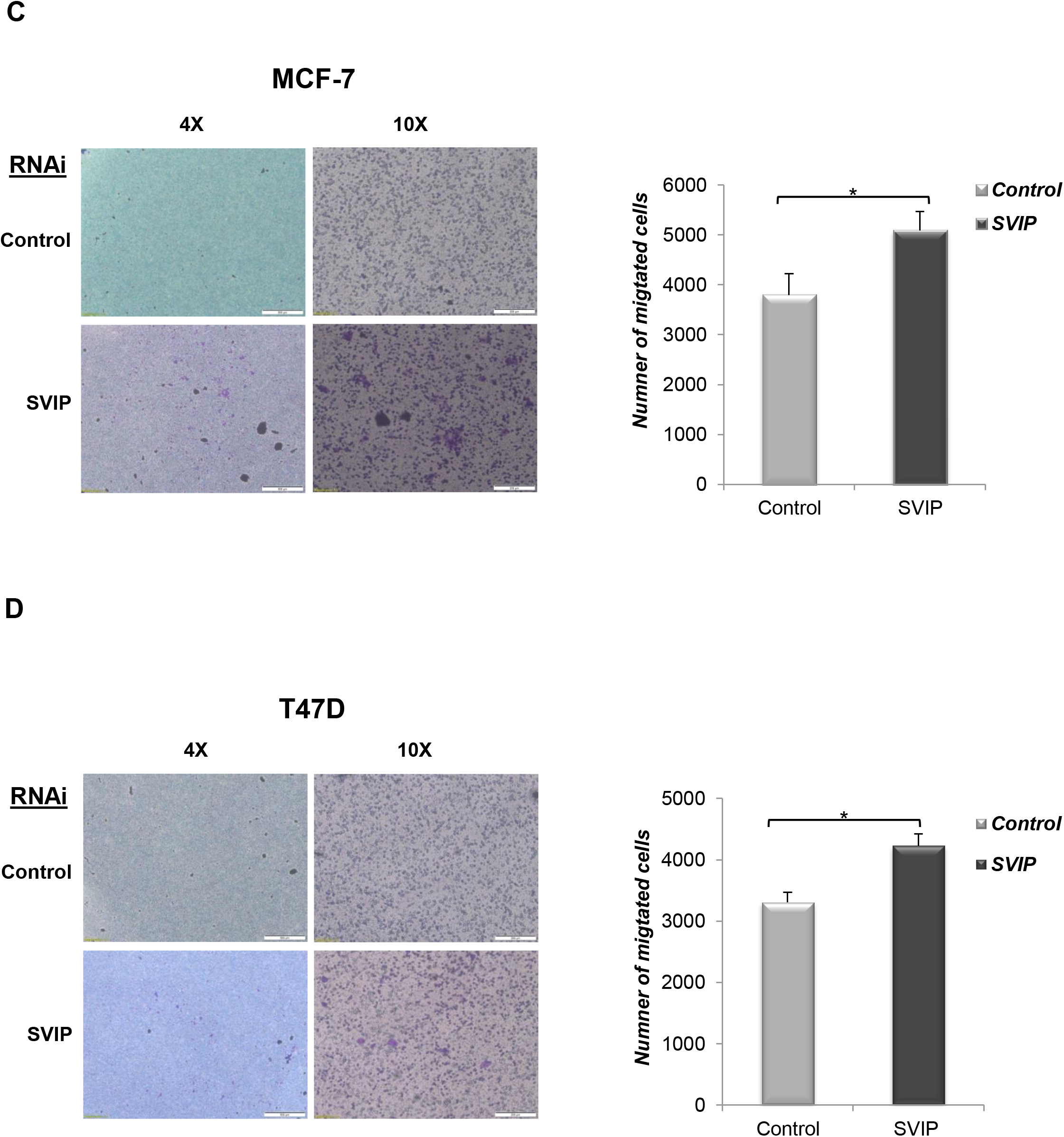
The role of SVIP in breast cancer tumorigenesis. Wound healing assay was performed in 0, 24, 48 h in transfected **A)** MCF7 and **B)** T47D cells. The analysis of wound closure % was determined using the ImageJ software. Two independent biological and three technical repeats per experiment were used. p-values were calculated with respect to control siRNA transfected cells by Student’s t-test (*p < 0.05). **C)** Transwell migration of MCF7 and **D)** T47D cell lines. Cells were allowed to migrate for 72h. Pictures were taken at ×4 and ×10 magnification. Three independent experiments were analyzed with student’s t-test (*p < 0.05). Error bars are presented as standard deviations

## 4. Discussion

The endoplasmic reticulum is involved in multiple cellular processes, including protein and lipid biosynthesis, protein folding and transport, and calcium hemeostasis. Nutrient deficiency, oxidative stress, high metabolic demand and calcium imbalance in the tumor microenvironment disrupt endoplasmic reticulum homeostasis and cause excessive accumulation of misfolded/unfolded proteins, with resultant endoplasmic reticulum stress. The major degradation pathway that endoplasmic reticulum uses is the ERAD pathway facilitating proteasome-dependent degradation of unfolded or misfolded proteins4. ERAD is a multi-component and complex system comprising many proteins, including gp78, Sel1L, Hrd1, Derlin-1/2 (Derl1/2), p97/VCP functioning on translocation, ubiquitination, and proteasomal degradation of non-native proteins. Besides functioning as a downstream response to UPR, recent evidence showed that ERAD complex has UPR-independent functions and there is a crosstalk between UPR and ERAD (Hwang and Qi et al., 2018).

The first indentified endogenous ERAD inhibitor SVIP is a multifunctional protein that has also been associated with autophagy, lysosomal dynamic stability, VLDL (very low-density lipoprotein) trafficking and secretion (Wang et al., 2011; Tiwari et al., 2016; Johnson et al., 2021). Furthermore, its gradually increasing expression in the developing nervous tissues, together with structural studies have been implicated that SVIP is a novel compact myelin protein (Wu et al., 2013; Ilhan et al., 2022). Very recently, SVIP was reported to be expressed in the adrenal gland at a consistently high level during postnatal development, suggesting that SVIP may function throughout the developmental process of the adrenal gland. Hence, modulation of SVIP expression was reported to alter not only the expression levels of steroidogenic genes and hormone output in adrenal cortex cells but also the expression of genes required for *de novo* cholesterol biosynthesis, uptake, and trafficking (Ilhan et al., 2022).

SVIP has recently started to attract attention in cancer biology studies. It has been reported that SVIP undergoes DNA hypermethylation–associated silencing in some cancer cells, especially in head and neck cancer. SVIP exhibited tumor suppressor features, and its recovery was found to be associated with increased ER stress and growth inhibition in SVIP-hypermethylated cancer cells. Interestingly, except for head and neck (50%), esophageal (23%), cervical (21%) cancers, and hematological malignancies (14%), particularly B cell lymphoma (33%), the SVIP promoter CpG island was most often found to be unmethylated in the other cancer types including prostate and breast cancers (Llinàs-Arias et al., 2019). SVIP has also been identified as androgen-regulated gene in prostate cancer and glioma (Erzurumlu and Ballar, 2017; Bao et al., 2017). It has been proved that SVIP is down-regulated, but other ERAD components and androgen receptor (AR) are upregulated in glioma and androgen-dependent prostate cancer cell lines with R1881 treatment. Furthermore, the regulation of the levels of SVIP and ERAD components leads to enhanced ERAD proteolytic activity, which was found to be related to prostate tumorigenesis. Similarly, the decreased SVIP expression, as well as increased AR expression, in glioma tissues correlated with gliomas progressing from low to high grades. Interestingly, the suppression of SVIP by AR was associated with decreased p53 expression, and overexpression of SVIP increased cell death only in p53wt glioma cell lines suggesting the downregulation of SVIP.

Apart from prostate cancer, breast cancer is another hormone-dependent cancer with a high incidence. Hormone receptor-positive breast and prostate cancers share several similarities, one of which is their dependence on the respective male and female hormones for their continued growth. While the estrogen and androgen promote the growth of these cancers, their deficiency results in low differentiation, increased proliferation rate, and unresponsiveness to the treatment (Ulm et al., 2019). As SVIP was identified as androgen-dependent gene and reported to be associated with tumorigenesis of both prostate cancer and glioma, we have investigated the SVIP expression in various cancers, with a particular focus on breast cancer due to its estrogen dependency.

First, the differential SVIP expression was observed in most human cancers through multi-omics data analysis. Our results revealed that SVIP expression was upregulated in many cancer types including adrenal, breast, and prostate cancers (**Fig. 1A**). Importantly, our data showed that SVIP expression was increased in all four intrinsic molecular subtypes of breast cancer [HER2+, Triple Negative (Basal Like), Luminal A, and Luminal B] compared to normal tissues using the GEPIA database (**Fig. S1**). On the other hand SVIP expression was significantly decreased especially in head and neck squamous cell carcinoma (**Fig. 1B**). These results are in line with a study by Llinàs-Arias et al. suggesting SVIP expression is upregulated in breast cancer, prostate cancer, lung cancer, and skin cancer while significantly decreased in head and neck tumors (Llinàs-Arias et al., 2019).

Epigenetic modifications, such as DNA methylation, have significant effect on tumorigenesis development and malignant transformation (Zhang et al., 2022). DNA methylation plays an important role in the progression and prognosis of breast cancer patients. Also, it has an important regulatory role in gene expression in cancer cells (Wu et al., 2021; Zhang et al., 2022). Previously, the downregulation of SVIP in some cancers, such as head and neck tumors, was shown to be associated with hypermethylation of SVIP promoter CpG island (Llinàs-Arias et al., 2019). Here, epigenetic-associated transcriptional regulation of SVIP was investigated in breast tumors, and the promoter methylation level of SVIP in breast cancer was found to be significantly lower than in normal tissues, and SVIP mRNA expression was negatively correlated with SVIP methylation (**Fig. 2A**). Consistently, the most common type of genomic alteration was high mRNA expression in all breast cancer types (**Fig. 2B**).

In addition to analysis of different online platforms confirming that the SVIP expression level is significantly higher in breast cancer than in normal tissues, our immunoblotting data also confirms that SVIP protein levels are significantly higher in breast cancer cell lines (MCF7, T47D, ZR75-1, BT-474, SK-BR-3 and MDA-MB-231) compared to human mammary epithelial cells (MCF-10A) (**Fig. 4A**). When the expression of key gp78-mediated ERAD pathway was evaluated we found that p97/VCP, Derlin1 and gp78 expression was higher only in MCF7 and BT474 cells. On the other hand, the main ERAD ubiquitin ligase Hrd1 had expression pattern similar to SVIP. This may be dues to the regulatory effect of both SVIP and Hrd1 on either the expression level or functionality of gp78. While SVIP inhibits gp78-mediated ERAD through removal of the p97/VCP and Derlin from gp78 complex, Hrd1 has been shown to targets gp78 for proteasomal degradation independent of the ubiquitin ligase activity of gp78 (Ballar et al., 2007; Shmueli et al., 2009; Ballar et al., 2010) Another possible reason may be the enhanced endoplasmic reticulum stress conditions in breast cancer and upregulation of Hrd1 and SVIP expressions by endoplasmic reticulum stress (Kikkert et al., 2004; Ballar et al., 2007).

It has been shown that endoplasmic reticulum stress impacts inflammatory and tumor microenvironment-induced immune suppression. Moreover, cancer cells transmit endoplasmic reticulum stress to immune cells recruited to inflammatory tissues (Zhou et al., 2022; Lou et al., 2023). Tumor-infiltrating immune cells play a key role against cancer treatment efficacy, an independent predictor of survival, prognosis and metastasis (Li et al., 2020; Cui et al., 2021). In recent years, the role of immune infiltrate cells in prognosis and response to treatments has become increasingly important also in breast cancer patients (Dieci et al., 2021). The immune cell infiltration in breast cancer is included different cellular subtypes like CD4+ and CD8+ T cells, B cells, monocytes, macrophages, dendritic cells, and natural killer cells (Goff et al., 2021). When the correlation of the SVIP expression with the tumor immune infiltration levels was investigated in breast cancer via TIMER, the results indicated that SVIP expression levels were positively associated with tumor purity (rho= 0.151, *p* = 1.60e-06) in breast cancer (Suppl Fig. 4). SVIP expression level had significant positive correlations with the infiltration level of CD8+ T cells (rho= 0.234, *p* = 8.11e-14), CD4+ T cells (rho=0.388, *p* =5.65e-37), B cells (rho= 0.121, *p* =1.29e-04) and Myeloid dendritic cells (rho= 0.156, *p* = 7.85e-07). However, there was a significantly negative correlation with the infiltration level of macrophage cells (rho= −0.143, *p* = 6.04e-06) and no significant correlation between the expression level of SVIP with the infiltration level of neutrophil (rho=0.045, *p* = 1.55e-01). Collectively, these results strongly exhibited that SVIP may also play an important role in immunotherapy using immune infiltrating cells in breast cancers.

Moreover, to investigate the impact of SVIP on cell proliferation and migration Real Time Cell Analyzer, wound healing and Transwell Boyden chamber analysis were performed in MCF7 and T47D cells. Previously, it has been reported that p53 wild type cell lines may be responsive to SVIP depletion by siRNA, thereby leading to increased cell proliferation of glioma cells; however, p53 mutant cell lines may not (Bao et al., 2017). On the other hand, the migration of both cell lines were affected by SVIP silencing, suggesting that breast cancer cells with lower SVIP expression might have higher migration ability (**Fig. 6A-D**). Interestingly, we found that SVIP expression is higher in primary breast tumors and lower in breast metastatic tumors compared to normal tissues (**Fig. 3C**). Consistent with these findings, breast cancer patients with lower SVIP expression exhibited a lower probability of survival compared to the patients with overexpressed SVIP (**Fig. 3D,E**). In brief, those results suggest that SVIP may be a prognostic biomarker for breast cancer.

In conclusion, as SVIP is an important regulator of proteostasis, its higher expression in breast cancer cell lines compared to normal breast cells as well as in breast cancer than in normal tissues indicates that SVIP might have a role in breast tumorigenesis. Furthermore, enhanced migration ability of SVIP silenced breast cancer cell lines, its higher expression in primary breast tumors but lower expression in breast metastatic tumors, which is well correlated with a lower probability of survival of breast cancer patients with lower SVIP expression compared to the patients with overexpressed SVIP, suggest that SVIP may be an important biomarker for clinical prediction on breast cancer. This study provides the foundation for future studies focusing on determining the role of SVIP in breast tumorigenesis.

## Abbreviations

UPR: unfolded protein response
ERAD: ER-associated degradation
SVIP: small VCP/p97-interacting protein
p97/VCP: valosin-containing protein
TIMER: the tumor immune estimation resource database
GEPIA: the gene expression profiling interactive analysis
TCGA: the cancer genome atlas
GTEx: the genotype-tissue expression project
bc-GenExMiner v4.8: the breast cancer gene-expression miner v4.
mRNA: messenger RNA
ER: estrogen receptor
PR: progesterone receptor
HER-2: Human epidermal growth factor receptor-2
E2: 17β-estradiol.

## Declaration of Competing Interest

The authors declare that they have no known competing financial interests or personal relationships that could have appeared to influence the work reported in this paper.

## Acknowledgements

This study was funded by the grants from Ege University Office of Scientific Research Projects (TDK-2021-22791 to PBK). EAS was supported by The Scientific And Technological Research Council of Türkiye (TUBITAK)-2211/C National PhD Scholarship Program in the Priority Fields in Science and Technology.

We thank the Pharmaceutical Sciences Research Centre (FABAL, Ege University, Faculty of Pharmacy) for equipment support.

## Data availability

Data will be made available on request.

## SUPPLEMANTARY FIGURE LEGENDS

**Fig. S1**. SVIP expression in breast cancer different subtypes and normal tissues (GEPIA). In the SVIP expression levels analysis using GEPIA, the threshold included expression fold change (Log2FC) ≥ 1 between breast cancer subtype and normal tissues, * p < 0.01. T = tumor tissues. N = normal tissues, NUM: number.

**Fig. S2**. Correlation of SVIP mRNA expression and methylation (cBioPortal). (Spearman=-019, p=8.81e-7; Pearson=-0.10, p= 0.0128)

**Fig. S3. A)** UALCAN, GEPIA, and bc-GenExMiner v4.8 portals analysis of breast cancer samples from the TCGA and GTEx datasets **B)** The mRNA expression level of SVIP in breast cancer patients with ER (−) and ER (+) (****, p < 0.0001) (bc-GenExMiner software). **C)** The mRNA expression level of SVIP in breast cancer patients with PR (−) and PR (+) (****, p < 0.0001) (bc-GenExMiner software). **D)** The mRNA expression level of SVIP in breast cancer patients with HER2 (−) and HER2 (+) (***, p = 0.0004) (bc-GenExMiner software). **E)** The mRNA expression level of SVIP in breast cancer patients with p53 wild type and p53 mutated (****, p < 0.0001) (bc-GenExMiner software).

**Fig. S4**.. Correlation of SVIP expression with immune infiltration level in breast cancer (TIMER) (*p* < 0.05).

## Notes

### Competing Interest Statement

The authors have declared no competing interest.

