## Supplementary figures and images for "Differential expression and function of SVIP in breast cancer cell lines and *in silico* analysis of its expression and prognostic potential in human breast cancer"

### Supplemental Figures

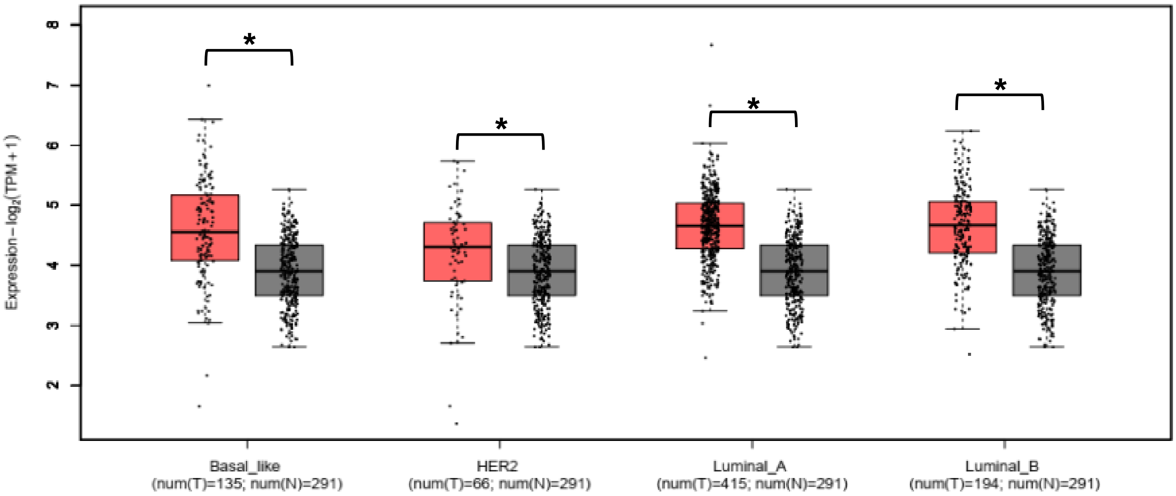

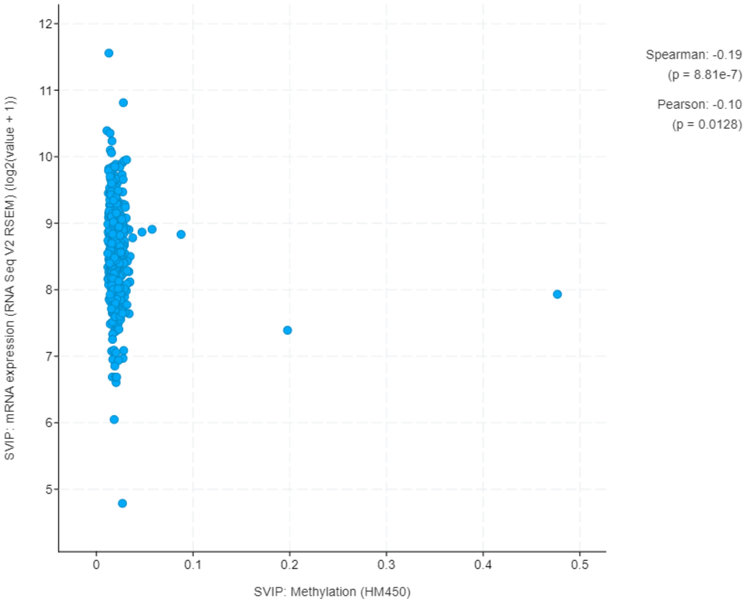

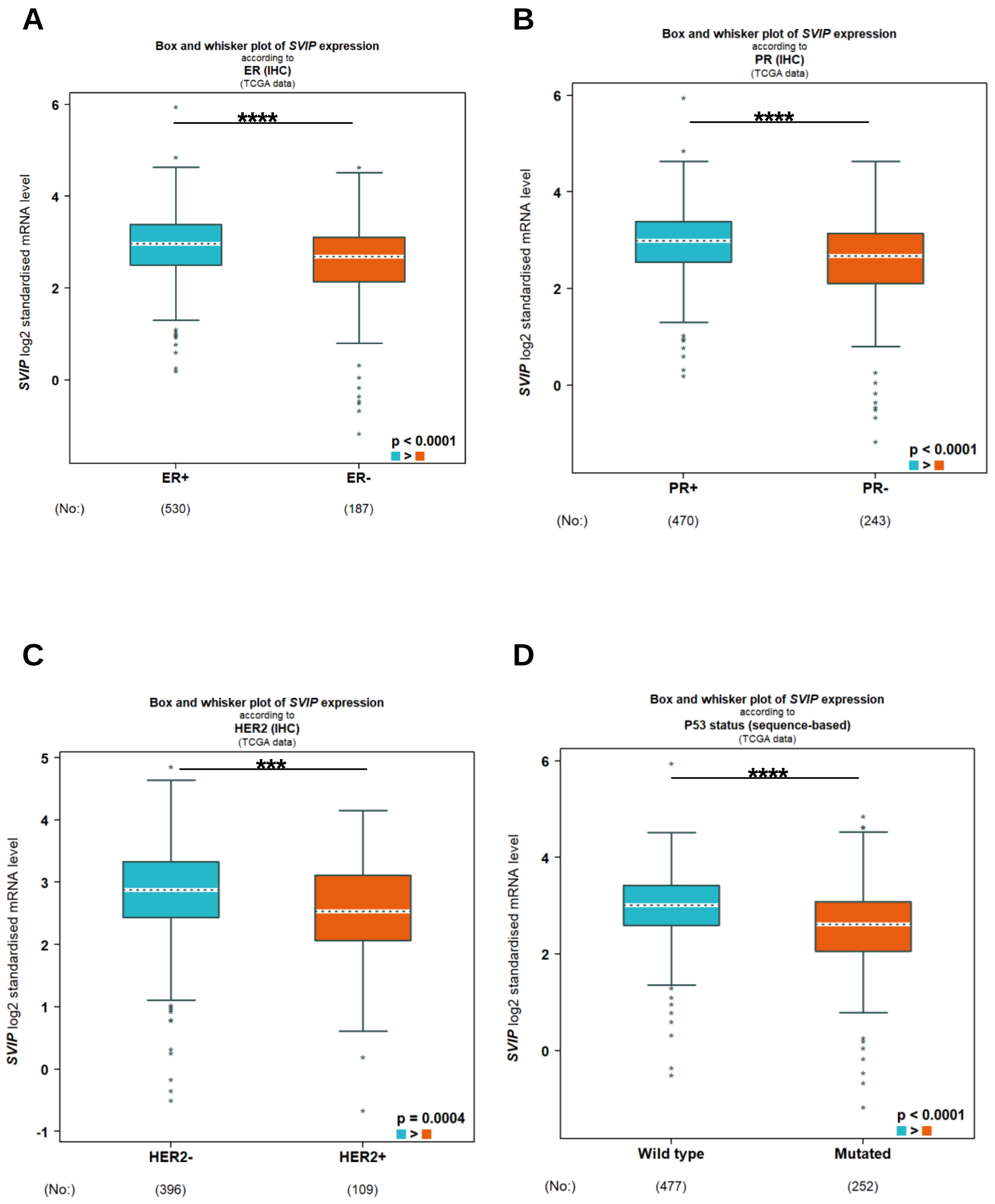

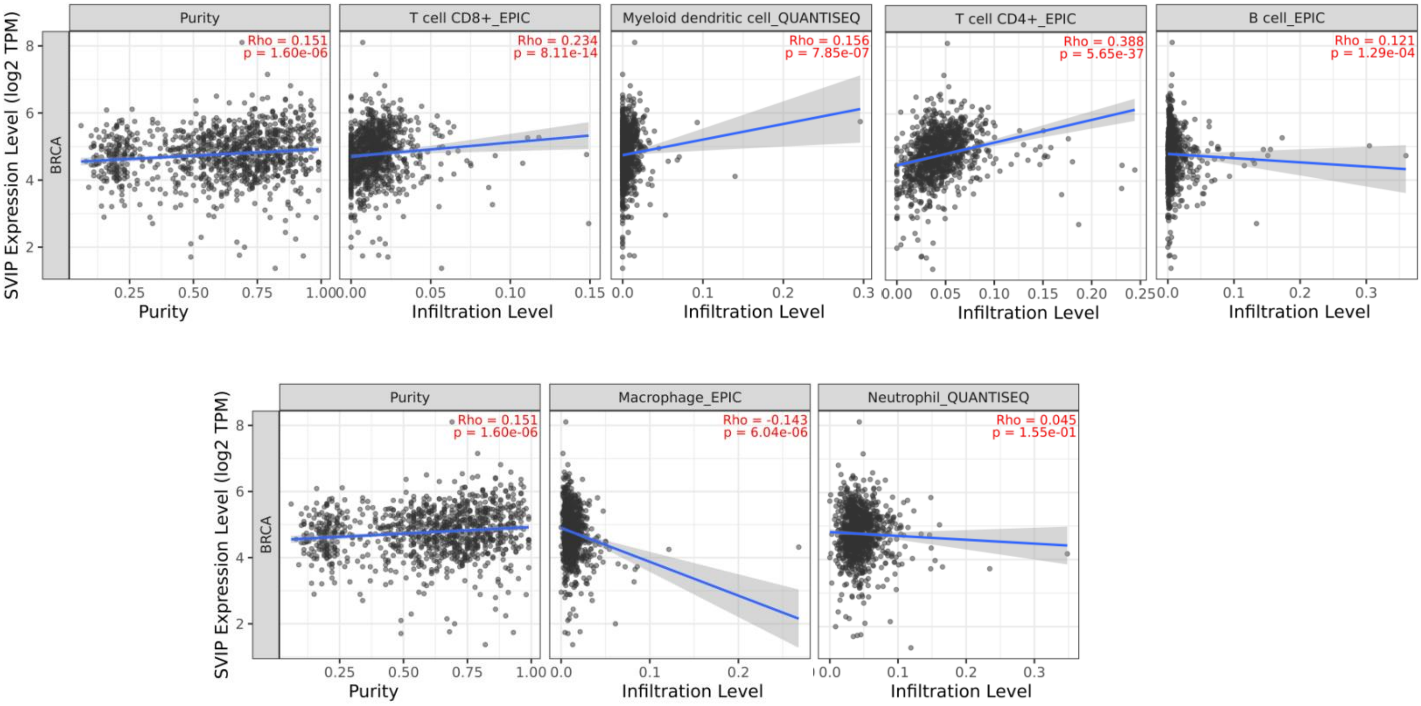
